# Lactate-Driven Heterogeneity of Immune Checkpoint Expression in Breast and Lung Cancer Cell Lines

**DOI:** 10.64898/2026.01.11.698903

**Authors:** Inigo San-Millan, Janel L. Martinez, Shivaun Lueke Pickard, Fred R. Hirsch, Christopher J. Rivard, George A. Brooks

## Abstract

Tumor-derived lactate is increasingly recognized as an immunosuppressive metabolite within the tumor microenvironment (TME), with emerging evidence highlighting its role beyond metabolism to include epigenetic and immune regulatory functions. While prior studies have primarily focused on individual immune checkpoints, most prominently PD-L1, it remains unclear whether lactate broadly coordinates the expression of multiple immune regulatory pathways across distinct tumor types, particularly in the context of chronic exposure mimicking glycolytic tumors. Here, we investigated the relationship between lactate-producing metabolism and immune checkpoint gene expression in four human cancer cell lines representing breast and lung cancer: MCF7 (estrogen receptor-positive breast), MDA-MB-231 (triple-negative breast), A549 (non-small cell lung), and H82 (small cell lung). By manipulating glucose availability and exposure duration to model acute (6 h) versus chronic (48 h) lactate production, and by pharmacologically inhibiting lactate dehydrogenase (LDH) with oxamate, we quantified extracellular lactate accumulation and assessed transcriptional responses of a panel of immune checkpoints (PD-L1, CD80, CD73, LGALS9, VISTA, PVR, CD47, FGL1, STING) and lactate-associated genes (MCT1, MCT4, LDHA, HCAR1) via qPCR. Chronic high-glucose conditions produced robust, LDH-dependent lactate accumulation and were associated with coordinated, lineage-specific remodeling of multiple checkpoint transcripts, whereas acute exposure induced minimal changes. MDA-MB-231 and A549 cells displayed striking but distinct checkpoint patterns under chronic lactate-producing conditions: MDA-MB-231 cells showed strong co-induction of PD-L1 and CD80, while A549 cells exhibited dominant CD80 induction with modest PD-L1 upregulation. H82 cells upregulated PD-L1 alongside CD73, LGALS9, CD47, and CD80, whereas MCF7 cells demonstrated more modest yet coordinated increases across several checkpoints. Chronic glucose exposure resulted in sustained, LDH-dependent lactate accumulation and coordinated induction of multiple immune checkpoint genes, with distinct lineage-specific patterns, e.g., robust PD-L1/CD80 upregulation in MDA-MB-231 versus CD80 dominance in A549. Unsupervised clustering and principal component analysis revealed that duration of glucose exposure, rather than acute glucose availability, was the primary axis of variation and that MCT4 and HCAR1 clustered with strongly induced checkpoints, consistent with a transcriptional program linking lactate export and sensing to immune regulation.

These findings support a model in which lactate acts as an upstream regulator of a broader immune escape program, potentially via mechanisms like lactylation and HCAR1 signaling. This work highlights the limitations of single-checkpoint blockade strategies in solid tumors and underscores the potential of targeting lactate metabolism to enhance immunotherapy efficacy in breast and lung cancers.

## INTRODUCTION

Immune checkpoint blockade has transformed the management of several cancers, yet most patients with solid tumors derive limited or no durable benefit, particularly in breast and lung malignancies (Pophali et al., 2024). Although antibodies targeting the PD-1/PD-L1 axis can trigger long-lasting remissions, primary and acquired resistance remain common, reflecting heterogeneous mechanisms that include poor T-cell infiltration, defective antigen presentation, and suppressive myeloid populations. Increasingly, dysregulated tumor metabolism is recognized as a critical, and targetable, determinant of immune escape(Cassim and Pouyssegur, 2019, Lei et al., 2020, Arner and Rathmell, 2023, Zhang et al., 2024, De Martino et al., 2024, Li et al., 2025, Wegiel et al., 2018, Reina-Campos et al., 2017, Lv et al., 2025, Cerezo and Rocchi, 2020, Leone and Powell, 2020).

Lactate, long considered a metabolic waste product of anaerobic glycolysis, is now recognized as a major signaling molecule (a “lacthormone”), a major gluconeogenic precursor and probably the preferred fuel for the body (Brooks, 2020, Brooks, 2018). In cancer, lactate accumulation, and not glucose oxidation, was what stuck Warburg in 1923 to posit cancer as an injury to cellular respiration (Warburg and Minami, 1923, Warburg et al., 1927, Warburg, 1925). Almost a decade ago, we developed the “lactagenesis hypothesis” by which we proposed that lactate was the reason and purpose of the Warburg effect, not for cellular bioenergetics but for signaling purpose for carcinogenesis where lactate is a key element in angiogenesis, immune escape, metastasis and self-sufficient metabolism of cancer cells (San-Millan and Brooks, 2017). Recently, we showed that lactate is an oncometabolite capable of regulating the expression of multiple driver genes in breast cancer (San-Millan et al., 2019, San-Millan et al., 2023).

Mechanistically, tumor-derived lactate influences gene expression and immune regulation through both cell surface signaling and epigenetic reprogramming. Recently, Di Zhang et al. elucidated a mechanism by which lactate directly regulates gene expression via histone lactylation, a post-translational modification in which lactate-derived lactyl groups are added to lysine residues on histones, promoting transcriptional activation (Zhang et al., 2019b, Zhang et al., 2024, Chen et al., 2025, Xu et al., 2024). Further, in lung cancer models, tumor cell-derived lactate promotes TAZ-dependent PD-L1 expression via GPR81 signaling, thereby dampening interferon-γ production and promoting T-cell dysfunction (Feng et al., 2017, Chen et al., 2025). These pathways position lactate as a nexus between metabolic rewiring and immune regulation within TME

Beyond direct effects on tumor cells, lactate exerts potent immunomodulatory actions. Elevated extracellular lactate suppresses proliferation, cytokine production, and cytotoxicity of CD8+ T cells and NK cells, while fostering regulatory T cells and myeloid-derived suppressor cells (Wegiel et al., 2018, Wang et al., 2021, Gu et al., 2025). Lactate also promotes M2-like polarization of tumor-associated macrophages and supports immune-suppressive macrophage phenotypes through both HIF-1α–dependent pathways and metabolic–epigenetic coupling (Colegio et al., 2014, Zhou et al., 2022, Xu et al., 2024, Zhang et al., 2022, Han et al.) Clinical and preclinical studies link high lactate or LDH activity to poor outcome and reduced T-cell infiltration, reinforcing the view that a lactate-rich TME is intrinsically hostile to antitumor immunity (Wegiel et al., 2018, Jedlička et al., 2022, Chen et al., 2024).

Clinically, Ddespite this rich mechanistic groundwork, most studies have focused on single checkpoints, particularly PD-L1, when exploring lactate’s impact on immune escape (Zhang et al., 2019a, Lei et al., 2020, Arner and Rathmell, 2023, De Martino et al., 2024). Yet tumors express diverse inhibitory ligands and receptors, including CD47, CD73, VISTA, LGALS9 (galectin-9), PVR, and CD80, that can independently or cooperatively suppress antitumor responses (Leone and Powell, 2020, Zhang et al., 2024, Lv et al., 2025).

Clinically, current immunotherapeutic strategies often rely on assessing PD-L1 expression to guide treatment decisions. This approach implicitly assumes that PD-L1 is the dominant inhibitory pathway in a given tumor (Pophali et al., 2024, Lei et al., 2020). When exploring lactate’s impact on immune escape PD-L1 is also the immune checkpoint with the highest emphasis on (Lei et al., 2020, Arner and Rathmell, 2023, De Martino et al., 2024). Yet tumors express diverse inhibitory ligands and receptors—including CD47, CD73, VISTA, LGALS9 (galectin-9), PVR, and CD80, that can independently or cooperatively suppress antitumor responses (Leone and Powell, 2020, Zhang et al., 2024, Lv et al., 2025). Failures of PD-1/PD-L1 inhibitors in many breast and lung cancers, together with emerging data linking lactate metabolism to resistance to anti-PD-1/PD-L1 therapy, suggest that alternative or compensatory checkpoints may predominate in lactate-high tumors (Zeng et al., 2024, Gu et al., 2025, Cerezo and Rocchi, 2020).

Here, we hypothesized that lactate-producing metabolism acts as an upstream coordinator of a multi-checkpoint immune escape program and that distinct tumor lineages adopt characteristic checkpoint signatures in response to chronic lactate-rich conditions. To address this, we quantified extracellular lactate accumulation and expression of a panel of immune checkpoints (PD-L1, CD80, CD73, LGALS9, VISTA, PVR, CD47, FGL1, STING) and lactate-associated genes (MCT1, MCT4, LDHA, HCAR1) across four breast and lung cancer cell lines (MCF7, MDA-MB-231, A549, H82) under acute versus chronic glucose exposure, with pharmacologic LDH inhibition to define LDH-dependent lactate production. Our goal was to confirm that sustained lactate-producing states are associated with coordinated remodeling of multiple immune checkpoints in a lineage-specific manner, with potential implications for immunotherapy resistance in lactate-rich breast and lung cancers.

## Methods

### 2.1 Cancer Cell Lines

Four human cancer cell lines were studied: MCF7 (estrogen receptor-positive breast adenocarcinoma), MDA-MB-231 (triple-negative breast cancer), A549 (non-small cell lung adenocarcinoma), and NCI-H82 (small cell lung carcinoma). Cells were obtained from the ATCC (Manassas, VA) and stored in liquid nitrogen until use.

Cells were grown on tissue culture-treated dishes (Genessee Scientific, San Diego, CA) in DMEM 1x media with 1 g/L glucose (10-014-CV, Corning, Manassas, VA), supplemented with 10% FBS and 1X penicillin/streptomycin. For experiments requiring high-glucose media, cells were switched to DMEM 1x with either 1 or 4.5 g/L glucose (10-013-CV, Corning). In these cases, the standard 10% FBS was replaced with 10% Nu-Serum (#51000, Corning Life Sciences, Glendale, AZ) to minimize initial lactate levels in the medium. All cell cultures were maintained at 37°C in a humidified incubator with 5% CO2 (Forma Isotemp 3530, ThermoFisher, Marietta, OH). Trypsin (0.25% trypsin, 25-053-CI, Corning) was used for subculturing and harvesting.

### 2.2 LDH Inhibition with Oxamate and Extracellular Lactate Quantification

To establish a lactate-rich extracellular environment independently of endogenous LDH-driven lactate production, selected experiments were performed in the presence of the LDH inhibitor sodium oxamate with or without the addition of exogenous L-lactate at the start of the incubation period. This approach allowed extracellular lactate availability to be experimentally controlled (“clamped”) under conditions of suppressed glycolytic lactate production. Total extracellular lactate after 48 h reflects the maintained lactate concentration in the conditioned medium, whereas net lactate values were calculated by subtracting the amount of lactate initially added to the medium, thereby providing an estimate of additional lactate accumulation or consumption during the incubation period.

#### Replication and Data Presentation

All conditions were performed in biological replicates (n=3 unless otherwise noted), and experiments were repeated independently at least twice. Lactate was quantified using a commercial assay kit (e.g., L-Lactate Assay Kit, Sigma-Aldrich, St. Louis, MO) following the manufacturer’s instructions. Data are presented as mean values ± standard error of the mean (SEM), with statistical significance determined by two-tailed t-tests (p<0.05).

### 2.3 qPCR Analysis

Following treatments, cells were trypsinized, resuspended in media, and pelleted by low-speed centrifugation. Total RNA was extracted from cell pellets using the RNeasy Plus Mini Kit (#74134, Qiagen, Hilden, Germany) per the manufacturer’s protocol. RNA concentration was measured using a Nanodrop 2000 spectrophotometer (Thermo Scientific, Waltham, MA). Complementary DNA (cDNA) was synthesized from 500 ng RNA per reaction using the iScript cDNA Synthesis Kit (#1708891, Bio-Rad, Hercules, CA).

Quantitative PCR (qPCR) was performed using probe-based primer sets and master mix (Prime Time, Integrated DNA Technologies, Coralville, IA) on a CFX96 real-time platform (Bio-Rad). Data were analyzed using Maestro software (Bio-Rad) and compared to a standard dilution curve generated from cell line cDNA. Housekeeping genes for normalization were selected using the Human Reference Gene Panel (Bio-Rad), with TBP and HPRT1 chosen based on stability across conditions.

#### Gene Expression Measurement

The qPCR panel targeted:

- Immune checkpoint ligands/receptors: PD-L1, CD47, CD73, CD80, FGL1, LGALS9, PVR, VISTA, and STING.
- Metabolic and lactate-associated genes: MCT1 (SLC16A1) and MCT4 (SLC16A3) (proton-coupled lactate transporters), LDHA (lactate dehydrogenase A, converting pyruvate to lactate), and HCAR1 (GPR81, G-protein-coupled lactate receptor).

Relative expression levels were calculated using the ΔΔCt method. Each gene’s expression in the −Glu 6 h condition (for each cell line) was used as the calibrator (set to 1.0), enabling fold-change calculations relative to this no-glucose baseline to isolate glucose/lactate-induced effects over time.

#### Data Visualization and Clustering

Fold-change expression data were compiled for all genes, cell lines, and time points. For visualization, a heatmap with hierarchical clustering was generated. Data were log_2_-transformed (fold-change of 1 = 0; >1 = positive/red; <1 = negative/blue) and centered at 0. Average linkage hierarchical clustering with a correlation distance metric was applied to group genes and conditions unbiasedly, using Python’s SciPy and Seaborn libraries.

For primary analysis, clustering focused on the 48 h time point to capture pronounced differences after prolonged lactate exposure. Additional clustering included 6 h data to assess early versus late responses. Dendrograms inferred relationships, such as cell line clustering by checkpoint profiles or gene grouping by patterns across models.

#### Statistical Analysis

As an exploratory study, changes (e.g., “upregulated” for >1.5-fold; “downregulated” for <0.67-fold) refer to consistent fold-changes across replicates. Where applicable, two-tailed t-tests compared gene expression between conditions (p<0.05 significant). Clustering robustness was evaluated by cophenetic correlation. All analyses were performed in Python or R for reproducibility.

## Results

### Chronic lactate-producing conditions across breast and lung cancer cell lines

To define the metabolic context for immune checkpoint regulation, extracellular lactate concentrations were quantified in four human cancer cell lines (MCF7, MDA-MB-231, A549, and H82) under glucose-deprived and high-glucose conditions at 6 h and 48 h. After 48 h of incubation in high-glucose medium (4.5 g/L), all cell lines accumulated substantial extracellular lactate, consistent with establishment of a chronic lactate-rich environment analogous to glycolytic tumors in vivo (Figure 2A). Absolute lactate levels differed between cell lines, reflecting known variation in glycolytic capacity and lactate export among breast and lung cancer subtypes.

**Figure 1.**
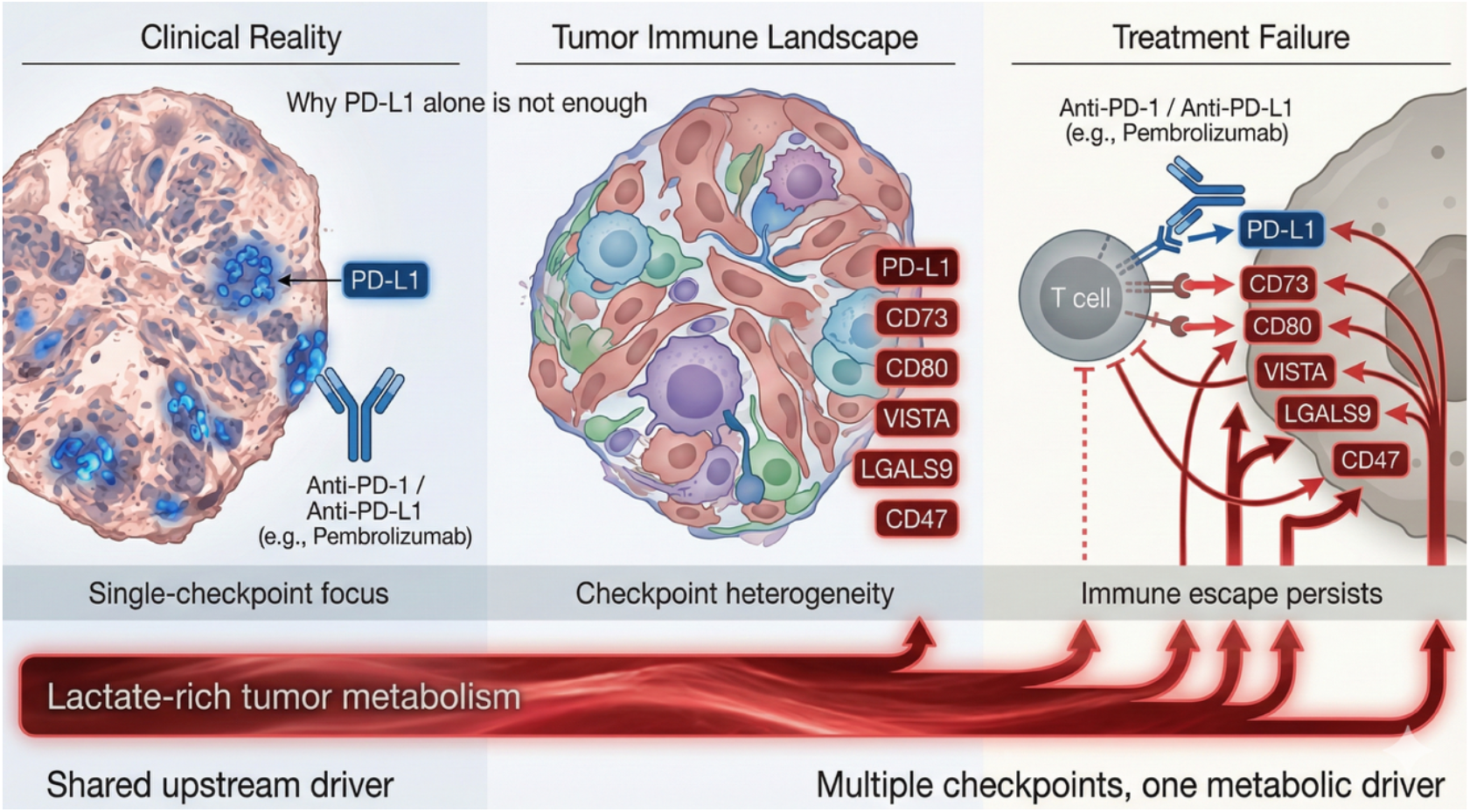
Conceptual framework linking lactate-rich tumor metabolism to multicheckpoint immune escape. Schematic illustration of the limitations of single-checkpoint–focused immunotherapy in the context of lactate-rich tumor metabolism. Left: clinical strategies often emphasize PD-L1 expression as the dominant immune escape mechanism and therapeutic target. Middle: tumors exhibit heterogeneous immune checkpoint landscapes, with coordinated expression of multiple inhibitory pathways including CD73, CD80, VISTA, LGALS9, and CD47. Right: blockade of the PD-1/PD-L1 axis alone may be insufficient to restore antitumor immunity when alternative checkpoints remain active. A shared lactate-rich metabolic state is proposed as an upstream condition associated with coordinated multicheckpoint regulation and persistent immune escape.

**Figure 2.**
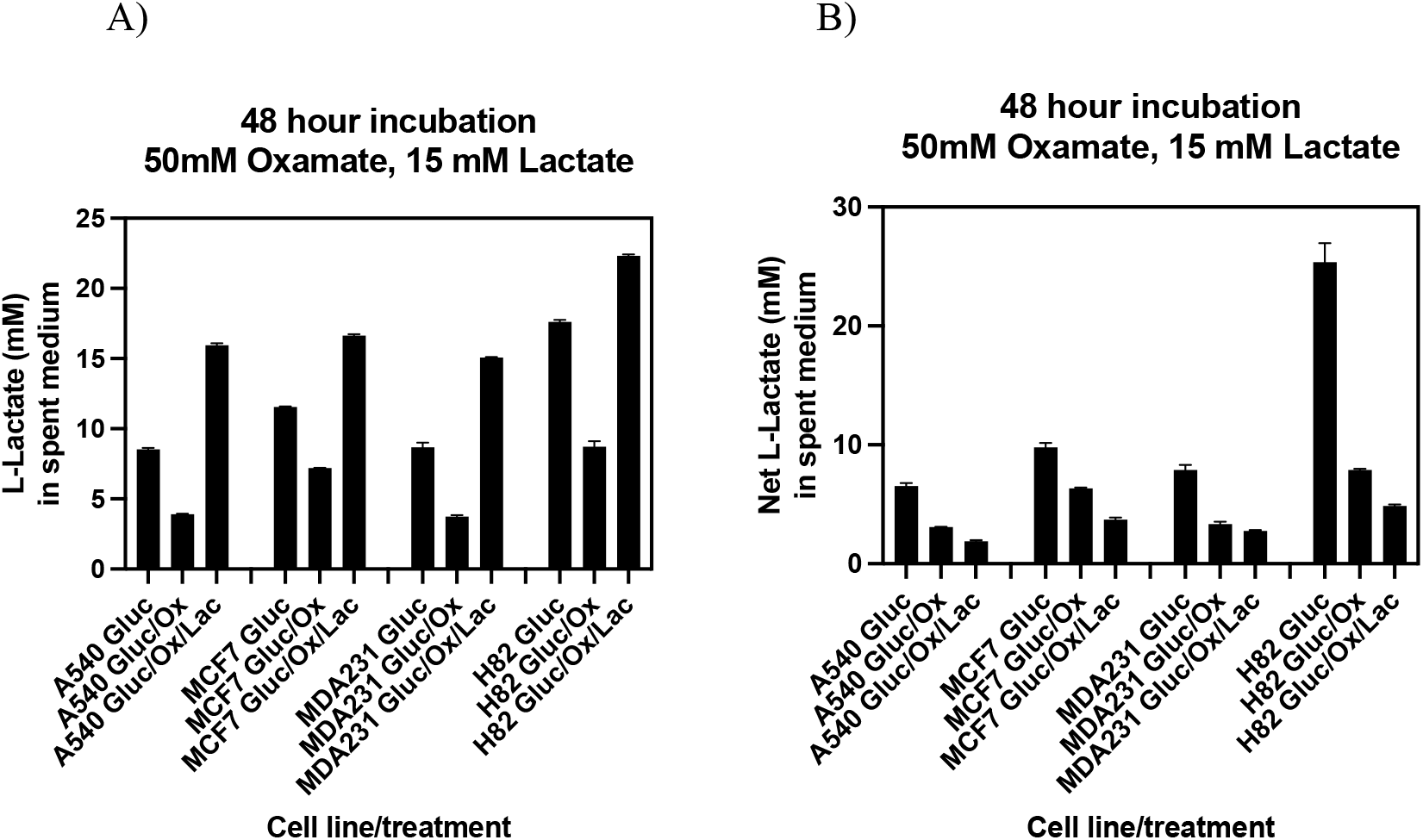
Extracellular lactate clamp under LDH inhibition distinguishes lactate availability from endogenous production. **(A)** Total extracellular L-lactate concentrations measured in spent medium from MCF7, MDA-MB-231, A549, and H82 cancer cell lines after 48 h incubation under glucose conditions, with or without lactate dehydrogenase (LDH) inhibition by oxamate (50 mM). Addition of exogenous L-lactate (15 mM) in the presence of LDH inhibition was used to establish an extracellular lactate clamp, maintaining high lactate availability independently of endogenous glycolytic lactate production. LDH inhibition markedly reduced lactate accumulation across all cell lines, confirming effective suppression of LDH-dependent lactate production, while exogenous lactate restored extracellular lactate levels to those observed under high-glucose conditions. **(B)** Net extracellular L-lactate accumulation over 48 h, calculated by subtracting baseline lactate levels from corresponding treatment conditions. Despite maintenance of high extracellular lactate levels under extracellular lactate clamp conditions, net lactate accumulation remained markedly reduced during LDH inhibition, consistent with suppression of endogenous lactate production and ongoing cellular lactate uptake and utilization. Data are presented as mean ± SEM.

Pharmacologic inhibition of lactate dehydrogenase (LDH) with oxamate markedly reduced extracellular lactate accumulation across all cell lines, confirming effective suppression of endogenous, LDH-dependent lactate production under these conditions. Importantly, addition of exogenous lactate in the presence of LDH inhibition re-established a sustained lactate-rich extracellular environment across all cell lines, thereby establishing an extracellular lactate clamp that maintained high lactate availability independently of endogenous glycolytic flux (Figure 2A). This experimental design enabled interrogation of lactate-associated immune regulatory programs while decoupling lactate availability from lactate production.

To further isolate glucose-derived lactate production, a net lactate accumulation metric was calculated by subtracting lactate levels measured under glucose-deprived conditions from those measured under corresponding high-glucose conditions. Notably, despite maintenance of high extracellular lactate levels under extracellular lactate clamp conditions, net lactate accumulation remained markedly reduced during LDH inhibition (Figure 2B).. This pattern is consistent with suppressed endogenous lactate production and ongoing cellular lactate uptake and utilization during the incubation period.

### Minimal checkpoint remodeling under acute glucose exposure

Immune checkpoint gene expression was next profiled by qPCR under acute (6 h) and chronic (48 h) glucose exposure, with expression normalized to the −Glu 6 h condition for each gene and line. Across all four cell lines, acute high-glucose exposure (6 h) produced only modest changes in checkpoint expression, with most genes remaining within ±1.2-fold of baseline (Figure 3). This limited response suggests that short-term changes in glucose availability, and the early phase of lactate production, are insufficient to drive broad transcriptional remodeling of immune checkpoint programs.

**Figure 3.**
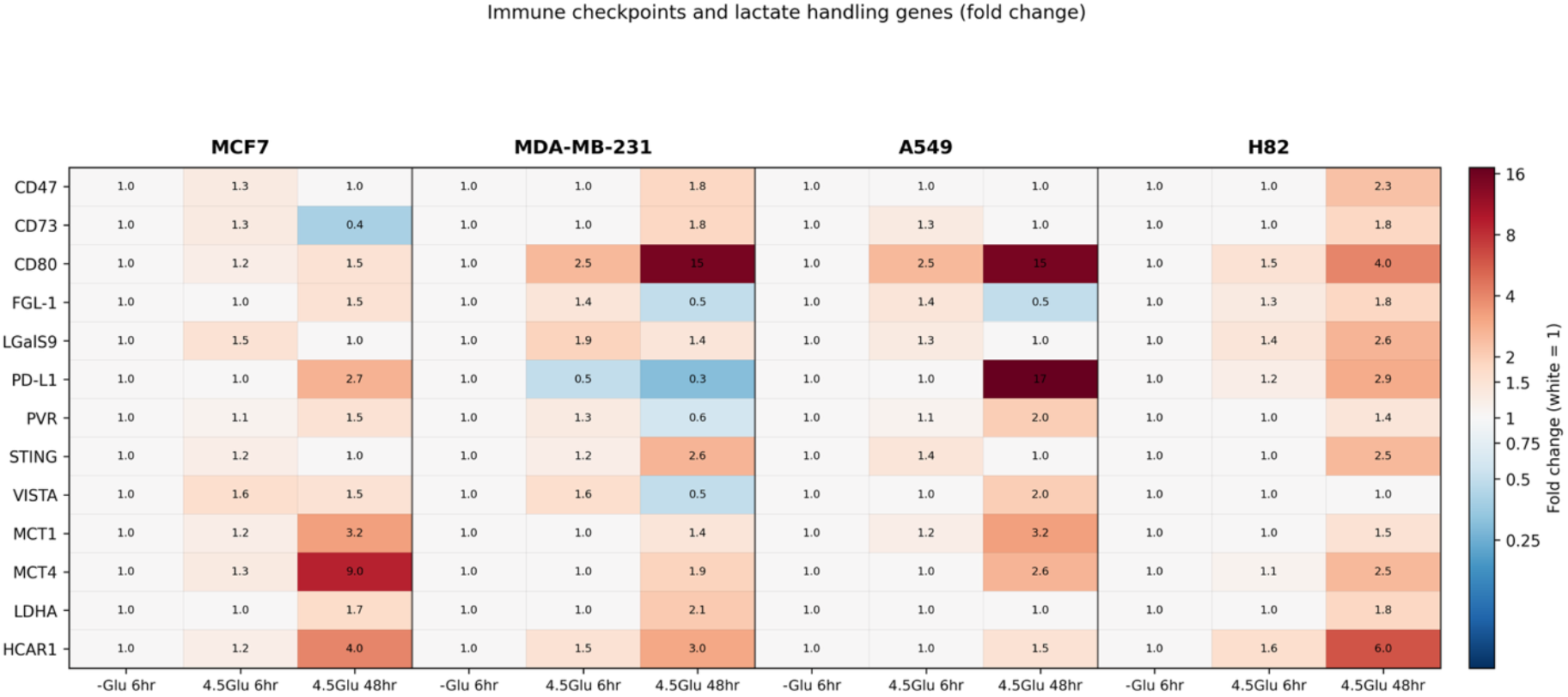
Chronic glucose exposure induces lineage-specific multicheckpoint transcriptional programs. Heatmap showing fold-change expression of immune checkpoint and lactate-handling genes in MCF7, MDA-MB-231, A549, and H82 cancer cell lines under glucose deprivation (−Glu, 6 h), acute glucose exposure (4.5 g/L, 6 h), and chronic glucose exposure (4.5 g/L, 48 h). Gene expression was quantified by qPCR and normalized to the −Glu 6 h condition for each gene and cell line (set to 1.0). Data are displayed as fold change, with white indicating no change, red indicating induction, and blue indicating repression. Chronic glucose exposure (48 h) resulted in coordinated but distinct immune checkpoint expression patterns across cell lines, whereas acute exposure produced minimal changes. Lactate transport and sensing genes (MCT1, MCT4, LDHA, HCAR1) exhibit parallel induction patterns consistent with adaptation to sustained lactate-rich metabolic states.

### Lineage-specific multi-checkpoint programs under chronic lactate-producing states

In contrast, chronic high-glucose exposure (48 h), which established a robust lactate-rich environment, was associated with pronounced and coordinated remodeling of multiple immune checkpoint transcripts in a lineage-dependent manner (Figure 2). In MCF7 cells, PD-L1, CD73, CD80, and LGALS9 were moderately but consistently induced (approximately 1.3–1.5-fold), whereas FGL1, PVR, and VISTA remained near baseline, suggesting a relatively low-amplitude but multi-component checkpoint response (Figure 3).

Checkpoint remodeling was more striking in MDA-MB-231 cells. PD-L1 and CD80 were strongly upregulated (approximately 15-fold) after 48 h in high glucose, representing the most robust transcriptional responses observed in this study, while CD73 showed induction of ∼1.8-fold and LGALS9 and CD47 were increased by ∼1.4–1.9-fold. FGL1 expression was reduced (∼0.5-fold), indicating that chronic lactate-producing conditions could simultaneously upregulate some inhibitory pathways and downregulate others. These features are consistent with an aggressive, PD-L1/CD80-centric immune escape module in triple-negative breast cancer under lactate-rich conditions (Figure 3)..

For cancer cells, A549 cells exhibited a distinct pattern. PD-L1 expression increased modestly (∼1.5-fold) at 48 h, whereas CD80 was strongly induced (∼15-fold), comparable in magnitude to MDA-MB-231. CD73 and LGALS9 showed modest increases (∼1.3–1.4-fold). FGL1 was downregulated (∼0.5-fold), similar to MDA-MB-231 (Figure 3).. These data suggest that in this NSCLC model, chronic lactate-producing states favor a CD80-dominant checkpoint program in which PD-L1 is not the primary inhibitory axis, with implications for PD-L1-based biomarker strategies.

H82 small-cell lung carcinoma cells adopted yet another checkpoint configuration under chronic high-glucose conditions. PD-L1 was induced by ∼2.9-fold, with concurrent upregulation of CD73 and CD47 (∼1.8–2.3-fold) and strong LGALS9 induction (∼2.6-fold). CD80 expression increased approximately 4-fold, indicating a significant, though less extreme, contribution of this checkpoint relative to MDA-MB-231 and A549 (Figure 3). Together, these patterns reveal that chronic lactate-producing conditions are linked to lineage-specific multi-checkpoint programs: high-amplitude PD-L1/CD80 in triple-negative breast cancer, CD80-dominant regulation in NSCLC, and a more distributed checkpoint architecture in SCLC and ER-positive breast cancer.

### Association of checkpoint remodeling with lactate transport and sensing

To determine whether immune checkpoint remodeling aligns with lactate handling capacity, expression of metabolic genes involved in lactate production, transport, and sensing was assessed in parallel. Across multiple cell lines, chronic high-glucose exposure induced MCT4 and HCAR1 expression, consistent with adaptation to sustained lactate export and signaling in a lactate-rich microenvironment. In contrast, LDHA expression changed modestly, suggesting that checkpoint remodeling is more tightly associated with lactate handling and sensing than with further upregulation of lactate-producing enzymes.

Unsupervised hierarchical clustering and principal component analysis were performed on normalized fold-change expression data to capture global patterns across genes, cell lines, and metabolic conditions (Figures 4 and 5). Samples segregated primarily according to the duration of glucose exposure, with 48 h conditions forming a distinct cluster separated from both 6 h glucose and glucose-deprived conditions across all cell lines, highlighting the dominant influence of chronic metabolic state. Notably, lactate-associated genes including MCT4 and HCAR1 showed strong co-loading with immune checkpoints such as PD-L1 and CD80, contributing prominently to principal components associated with prolonged high-glucose exposure.

**Figure 4.**
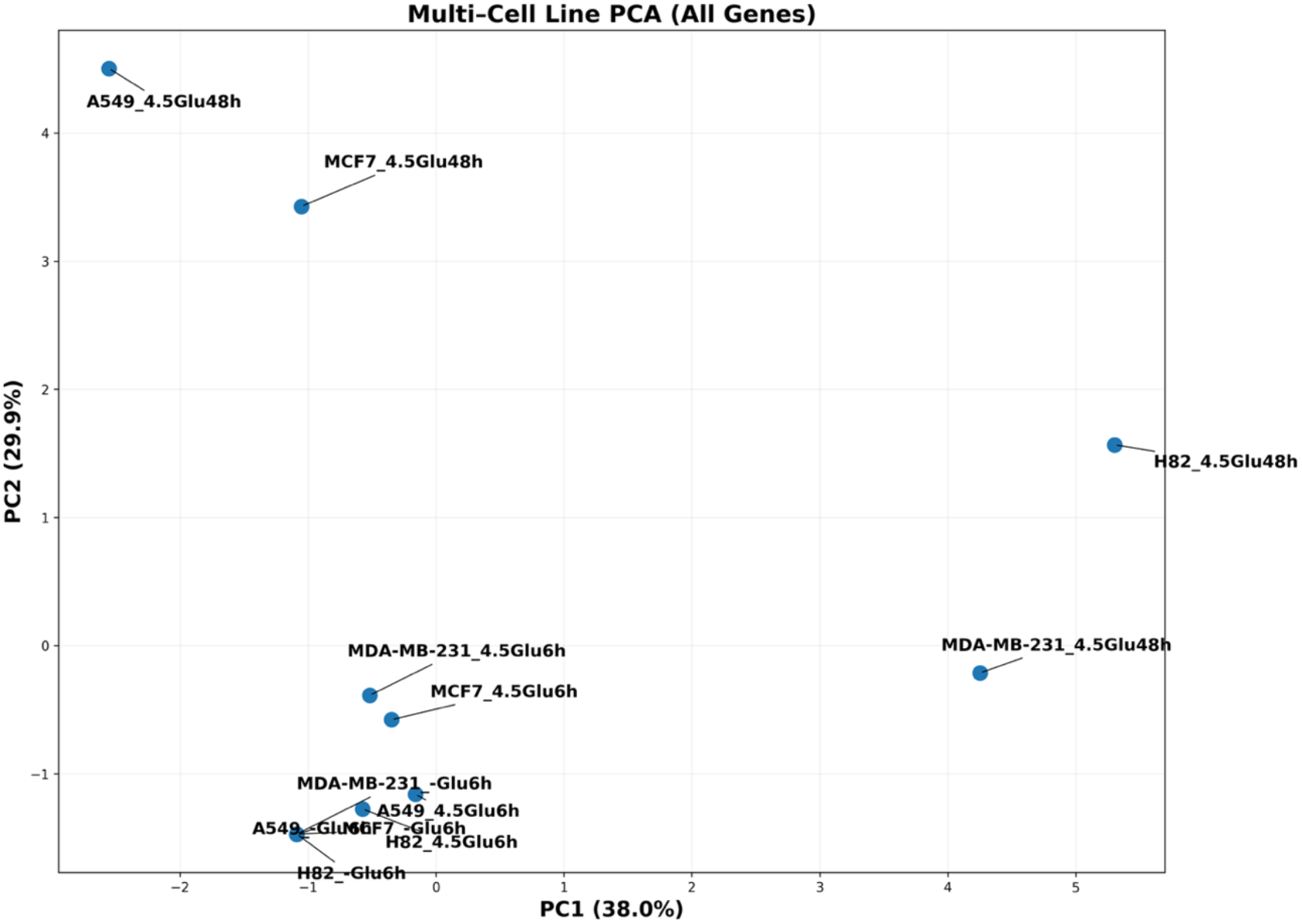
Principal component analysis reveals duration of metabolic exposure as the dominant driver of transcriptional variation. Principal component analysis (PCA) of normalized fold-change expression data for immune checkpoint and lactate-handling genes across four cancer cell lines (MCF7, MDA-MB-231, A549, and H82) under glucose deprivation (−Glu, 6 h), acute glucose exposure (4.5 g/L, 6 h), and chronic glucose exposure (4.5 g/L, 48 h). Each point represents a single cell line–condition pair. Samples segregate primarily along PC1 according to duration of glucose exposure, with chronic (48 h) conditions clearly separated from acute and glucose-deprived states across all cell lines. PC2 captures lineage-specific variation among tumor models. Percent variance explained by each principal component is indicated on the axes.

**Figure 5.**
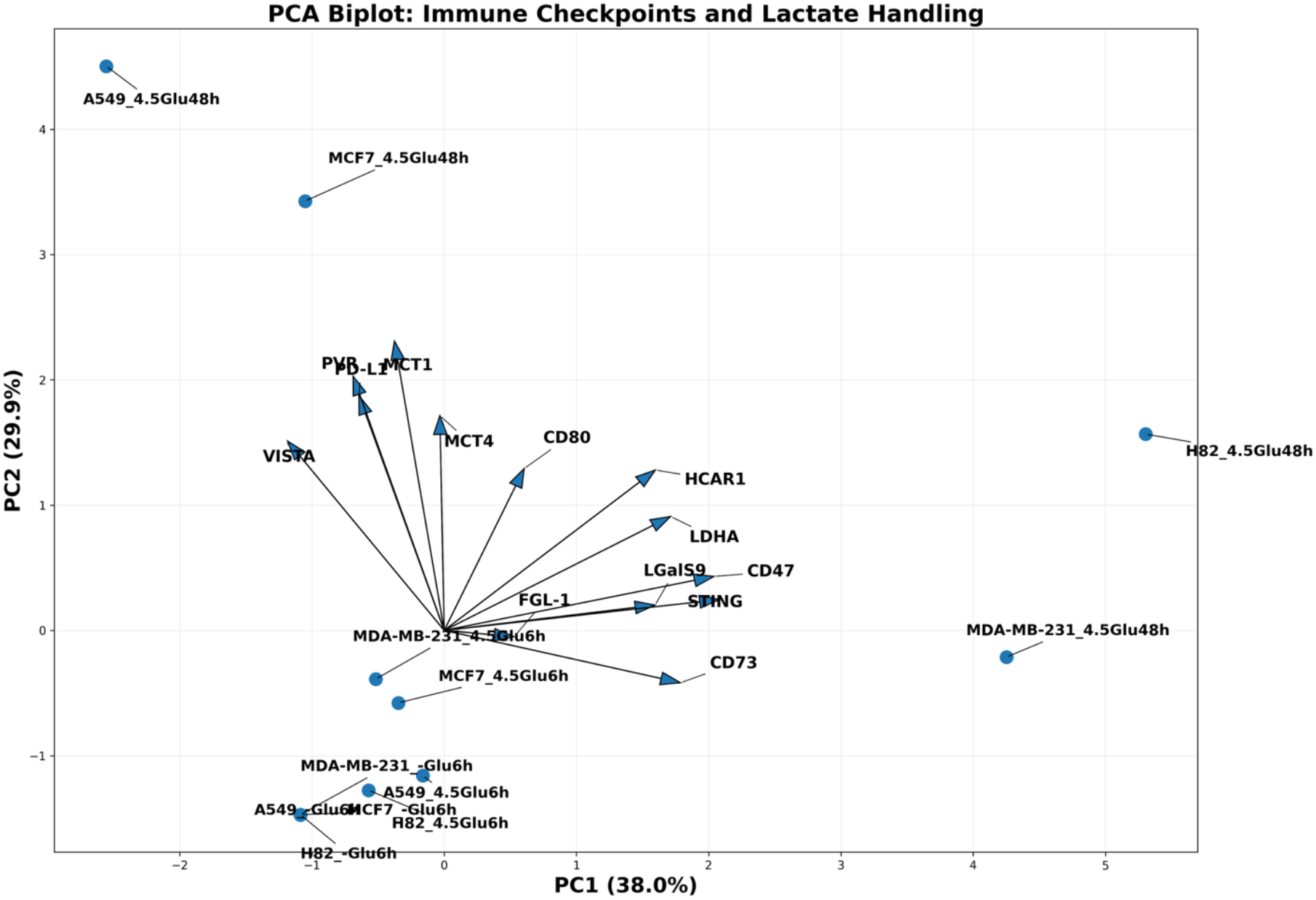
PCA biplot showing coordinated metabolic and immune checkpoint regulation under chronic glucose exposure. Biplot representation of the PCA shown in Figure 4, displaying both sample scores (points) and gene loading vectors (arrows). Arrows indicate the direction and relative contribution of individual genes to the principal components. Lactate handling and sensing genes, including MCT4 and HCAR1, together with selected glycolytic markers, co-load with immune checkpoints such as PD-L1 and CD80 along principal components associated with chronic glucose exposure (48 h). Samples exposed to prolonged high-glucose conditions cluster in the direction of these vectors, consistent with coordinated transcriptional remodeling of metabolic and immune programs under sustained lactate-producing states.

Together, these analyses demonstrate that sustained lactate-producing metabolic states drive coordinated, multi-checkpoint immune remodeling, with the dominant checkpoints differing by tumor lineage.

## Discussion

This work shows that chronic lactate-producing metabolic states are associated with coordinated remodeling of multiple immune checkpoint genes in breast and lung cancer cell lines, with the magnitude and pattern of checkpoint induction varying by tumor lineage. Acute glucose exposure produced minimal changes in checkpoint expression, whereas prolonged high-glucose conditions, which drove robust LDH-dependent lactate accumulation, were linked to pronounced upregulation of multiple checkpoint transcripts. These observations support a model in which sustained lactate-rich conditions, rather than transient nutrient availability, favor stable immune regulatory programs at the transcriptional level (Wang et al., 2021).

The observation that net lactate accumulation remains reduced under LDH inhibition, even when extracellular lactate is experimentally clamped at high levels, likely reflects continued lactate uptake and oxidation by cancer cells, highlighting the dynamic nature of lactate flux and reinforcing the distinction between lactate availability and lactate production.

From a translational perspective, current immunotherapy strategies frequently emphasize PD-L1 expression as a biomarker and therapeutic target. However, our data suggest that tumors may rely on alternative or compensatory inhibitory pathways, such as CD73, VISTA, LGALS9, or CD80, particularly under chronic lactate-rich conditions. In triple-negative MDA-MB-231 cells, PD-L1 and CD80 were strongly induced, while CD73, LGALS9, and CD47 showed more modest increases and FGL1 was downregulated, suggesting a high-amplitude, PD-L1/CD80-centric immune escape module. In A549 non-small cell lung cancer cells, CD80 induction was striking whereas PD-L1 upregulation was modest, pointing to a CD80-dominant strategy that may not be fully captured by PD-L1–based biomarker approaches. H82 small-cell lung carcinoma cells exhibited intermediate PD-L1 induction together with upregulation of CD73, LGALS9, CD47, and CD80, consistent with a more distributed multi-checkpoint architecture. MCF7 cells displayed coordinated but modest increases across several checkpoints, suggesting a lower-intensity but still multi-component program. This heterogeneity may help explain why PD-1/PD-L1 blockade is highly effective in some cancers yet fails in many solid tumors, including breast and lung, where immune suppression is distributed across multiple checkpoints and influenced by metabolic cues like lactate accumulation. Emerging evidence shows that lactate inhibits PD-1/PD-L1 interactions and promotes resistance to checkpoint inhibitors in lactate-high environments (Oh et al., 2025). Moreover, These lineage-specific patterns resonate with recent work indicating that tumor-intrinsic metabolic phenotypes govern distinct modes of immune escape (Chen et al., 2024). Combinatorial approaches targeting lactate production or sensing together with multi-checkpoint blockade may be needed to fully reverse immune escape in breast and lung cancers.

The clustering of immune checkpoints with lactate handling and sensing genes strengthens the link between lactate metabolism and immune regulation at the systems level. Chronic glucose exposure induced MCT4 and HCAR1 expression, and these genes clustered with strongly induced checkpoints such as PD-L1 and CD80 in hierarchical and principal component analyses. This configuration is consistent with prior mechanistic studies in which lactate–HCAR1/GPR81 signaling increased PD-L1 expression through cAMP/PKA–TAZ pathways (Feng et al., 2017) and with reports that lactate-driven histone lactylation can activate gene programs that promote tumor progression and immune suppression (Zhang et al., 2019b, Chen et al., 2025). Together with extensive literature showing that lactate suppresses effector T and NK cells while favoring regulatory and myeloid suppressor populations (Wang et al., 2021, Jedlička et al., 2022, Gu et al., 2025), these data support a unified view of lactate as a central coordinator of both metabolic and immune dimensions of the TME.

Targeting lactate metabolism and transport offers several potential points of intervention. Blockade of HCAR1/GPR81 signaling has been shown to decrease PD-L1 expression, impair tumor growth, and synergize with metabolic agents in vivo (Feng et al., 2017, Gu et al., 2025, Llibre et al., 2025), suggesting that disrupting lactate sensing may attenuate checkpoint upregulation at its source. Modulation of lactylation or related metabolic–epigenetic pathways in tumor and myeloid cells may further reprogram checkpoint expression and macrophage polarization (Chen et al., 2025, Jin et al., 2025), opening avenues to remodel the immunosuppressive TME. The lineage-specific checkpoint patterns described here provide a rationale for tailoring such interventions to tumor subtype, for example, combining LDH/HCAR1 blockade with CTLA-4–axis targeting in CD80-dominant NSCLC or with multi-checkpoint inhibition in TNBC with high PD-L1/CD80 expression.

A key limitation of this study is the absence of direct transcriptional measurements under LDH inhibition conditions, as well as reliance on in vitro models without functional immune assays or protein-level validation. Nevertheless, the strong temporal association between LDH-dependent lactate accumulation and immune checkpoint remodeling, coupled with clustering of lactate-handling genes like MCT4 and HCAR1 with induced checkpoints, supports a functional link between lactate-producing metabolism and immune regulation. Future studies incorporating genetic or pharmacologic disruption of lactate signaling pathways (e.g., HCAR1 antagonists), coupled with co-culture models or in vivo validation, will be essential to establish causality and explore lactylation’s role in these processes.

Future work should therefore integrate genetic and pharmacologic perturbations of lactate metabolism and signaling with co-culture or in vivo models that incorporate key immune effectors. For example, exogenous lactate titration studies combined with HCAR1 antagonists or lactylation modulators could test whether the multi-checkpoint transcriptional programs described here are lactate-dependent and receptor- or epigenetic-mediated. Co-cultures with T cells, NK cells, and macrophages, or syngeneic and humanized mouse models, could then link specific checkpoint patterns to functional outcomes such as T-cell activation, cytotoxicity, macrophage polarization, and response to checkpoint blockade in lactate-rich tumors. By connecting the lineage-specific checkpoint signatures observed here with mechanistic and functional readouts, it should be possible to refine lactate-targeted immunometabolic strategies and identify combination regimens best suited for distinct breast and lung cancer subtypes.

In summary, our findings support a model in which lactate acts as a central regulator of immune checkpoint expression across tumor types. By highlighting the heterogeneity and coordination of immune checkpoints in response to metabolic cues, this work underscores the need to move beyond single-checkpoint paradigms and to consider tumor metabolism as a key determinant of immunotherapy response in breast and lung cancers.

